# Development of humanized mouse and rat models with full-thickness human skin and autologous immune cells

**DOI:** 10.1101/2020.01.14.906255

**Authors:** Yash Agarwal, Cole Beatty, Sara Ho, Lance Thurlow, Antu Das, Samantha Kelly, Isabella Castronova, Rajeev Salunke, Shivkumar Biradar, Tseten Yeshi, Anthony Richardson, Moses Bility

**Author notes:** The National Institute of Health, which funds this work, “requires scientists to submit final peer-reviewed journal manuscripts that arise from NIH funds to the digital archive PubMed Central upon acceptance for publication.”. denotes authors contributed equally to the work (Yash Agarwal, Cole Beatty, Sara Ho). denotes corresponding author.

## Abstract

The human skin is a major barrier for host defense against many human pathogens, with several pathogens directly targeting the skin for replication and disease. The skin is also the primary route of infection for a myriad of vector-borne diseases; thus cutaneous immune cells play a major role in modulating transmission for such infectious diseases. Several human pathogens that target the skin as a major route of infection are unable to infect traditional rodent models or recapitulate the pathogenesis in humans. It is well established that differences exist in human skin and immune cell biology compared to rodent models. Therefore, rodent (mouse and rat) models that incorporate human skin and immune cells would addressed the above discussed technical gap, and enable *in vivo* mechanistic studies of human host-skin pathogen interactions, and support the development of novel therapeutics. Here, we introduce the novel human Skin and Immune System (hSIS)-humanized NOD-*scid* IL2Rg^null^ (NSG) mouse and Sprague-Dawley-Rag2^tm2hera^ Il2rg^tm1hera^ (SRG) rat models, co-engrafted with full-thickness human fetal skin, autologous fetal lymphoid tissues, and fetal liver-derived hematopoietic stem cells. hSIS-humanized rodents support the development of adult-like, full-thickness human skin and human lymphoid tissues, and support human immune cell development. Furthermore, the engrafted human skin supports Methicillin-resistant *Staphylococcus aureus* infection, demonstrating the utility of these humanized rodent models in studying human disease.

## Introduction

The human skin is a major target for infection or transmission of several infectious agents of global health significance ^1,2^. Several emerging pathogens, namely community-associated methicillin-resistant *Staphylococcus aureus* (CA-MRSA) and *Candida auris*, respectively, target the skin for infection and disease ^1,3-5^. Vector-borne infectious diseases such as Lyme disease, malaria, and dengue are transmitted via ticks and mosquito respectively, via inoculation in the skin. Recent evidence suggests that the cutaneous immune system plays a critical role in modulating the establishment of infections in the skin ^6^. In concurrence with the skin’s role as a major barrier for host defense against infectious agents, the cutaneous immune system plays a major role in initiating systemic immune response and abrogates pathogen replication and trafficking to other sites of replication ^7-10^. Furthermore, the cutaneous immune system provides an ideal target for inducing human immunity against various infectious agents via vaccination with novel vaccine technologies, such as skin-patch intradermal vaccines ^11,12^. The skin is also a major barrier against environmental insults, including ultraviolet radiation from the Sun; and it is well established that long-term overexposure to ultraviolet radiation induces cutaneous immune suppression ^13^.

*In vivo* models for studying pathogens that target the skin for infection or transmission primarily involve mice and rats ^3^. Said rodent models have improved mechanistic understanding of the transmission, replication, and pathogenesis of a myriad of human pathogens; however, significant differences exist between the skin and immune system (including the cutaneous immune system) of humans and rodents ^3,14^. Consequently, translational gaps can form between clinical outcomes and studies performed with conventional rodent models ^1^. To addressed the above discussed technical gaps, the severely immunodeficient NOD-*scid* IL2Rg^null^ (NSG) mouse model, which lacks mature T cells, B cells, and natural killer (NK) cells, along with defects in innate immunity, has been engrafted with various human cells and tissues (termed, humanized NSG mice) to enable investigation of human host-pathogen interactions and recapitulate clinical features, ^15-17^. Recently, a severely immunodeficient rat, termed, Sprague-Dawley-Rag2^tm2hera^ Il2rg^tm1hera^ (SRG) rat ^18,19^, with comparable immunodeficiency as in the NSG mouse model, was developed to support engraftment of human cells and tissues into a larger and longer-life span rodent model. To date, a humanized rodent model co-engrafted with full-thickness human skin and autologous lymphoid tissues and immune cells (including cutaneous immune cells) remains to be developed and established; albeit mouse and rat models engrafted with human skin, with or without allogeneic human peripheral blood mononuclear cells (PBMCs) (with human T cells the predominant immune cell subset that engraft) have been established ^20-23^.

Here, we employ the severely immunodeficient NSG mouse and SRG rat models to generate human Skin and Immune System (hSIS)-NSG mice and SRG rats for investigating human skin-associated infections and diseases. NSG mice and SRG rats were co-engrafted with full-thickness human fetal skin over the rodent skin-excised rib cage, autologous fetal lymphoid tissues under the renal capsule, along with autologous fetal liver-derived hematopoietic stem cells via intravenous injection. We report that the hSIS-humanized NSG mouse model supports the development of full-thickness human skin, autologous human immune cells, and human thymus and spleen tissues. The hSIS-humanized SRG rat model also supports the development of full-thickness human skin, thymus tissue and autologous human immune cells. We demonstrated that hSIS-humanized rodents support CA-MRSA infection and associated skin pathology to recapitulate clinical presentations in humans.

## Results

### The hSIS-humanized NSG mouse supports the development of full-thickness human skin, autologous lymphoid tissues (thymus and spleen), and human immune cells

We previously demonstrated that severely immunodeficient NSG mice support robust development of human lymphoid tissues (thymus and spleen) along with autologous immune cells ^24^; separate studies demonstrated that severely immunodeficient mice support the engraftment and development of human skin ^22^. The mouse skin microanatomy differs from human skin microanatomy by the absence of multi-layered epidermis, eccrine and apocrine glands, and the papillary, reticular and hypo-dermal regions of the dermal layer ^25^. Additionally, the microanatomy of human primary and secondary lymphoid tissues differs significantly from mouse lymphoid tissues, with significant differences in red pulp to white pulp ratio in the spleen and lobulation in the thymus ^24,26,27^. Here, we hypothesize that severely immunodeficient NSG mice will support co-engraftment of full-thickness fetal human skin and autologous fetal lymphoid tissues along with hematopoietic stem cells. Furthermore, we hypothesize that severely immunodeficient NSG mice will facilitate human skin and lymphoid tissues development, and enable systemic human immune cell reconstitution in transplanted human tissues and the blood (**Figure 1**). We processed human fetal spleen, thymus, and liver organs into ∼1 mm^2 tissues and isolated autologous human CD34^+^ hematopoietic stem cells from the fetal liver, and transplanted the tissues and hematopoietic stem cells into irradiated NSG mice (**Figure 1**). Full-thickness human fetal skin was processed via removal of excess fat tissues attached to the subcutaneous layer of the skin, then engrafted over the rib cage where mouse skin was previously excised (**Figure 1**). Gross analysis of human skin in hSIS-NSG mice beginning at 2 weeks post-transplantation demonstrates robust wound healing and maturation into adult-like human skin (10 weeks post-transplantation) (**Figure 2A**). Gross analysis of human spleen and thymus tissues development in hSIS-humanized NSG mice at 10 weeks post-transplantation demonstrated the development of human spleen and thymus tissues under the kidney capsule (**Figure 2B**) ^24^. In addition to supporting the development of human spleen and thymus tissues, the hSIS-humanized NSG mouse model supports reconstitution of the immunodeficient-murine lymph nodes and spleen (**Figure 2B**) ^24^. The human skin in hSIS-NSG mice also develops human skin appendages (hair) (**Supplementary Figure 1**). A limitation of the human skin in the hSIS-NSG mouse model is the development of dry skin and early signs of graft-versus-host disease in and around the human skin graft at 20 weeks post-transplantation, which is associated with rapid aging (**Figure 3**).

**Figure 1.**
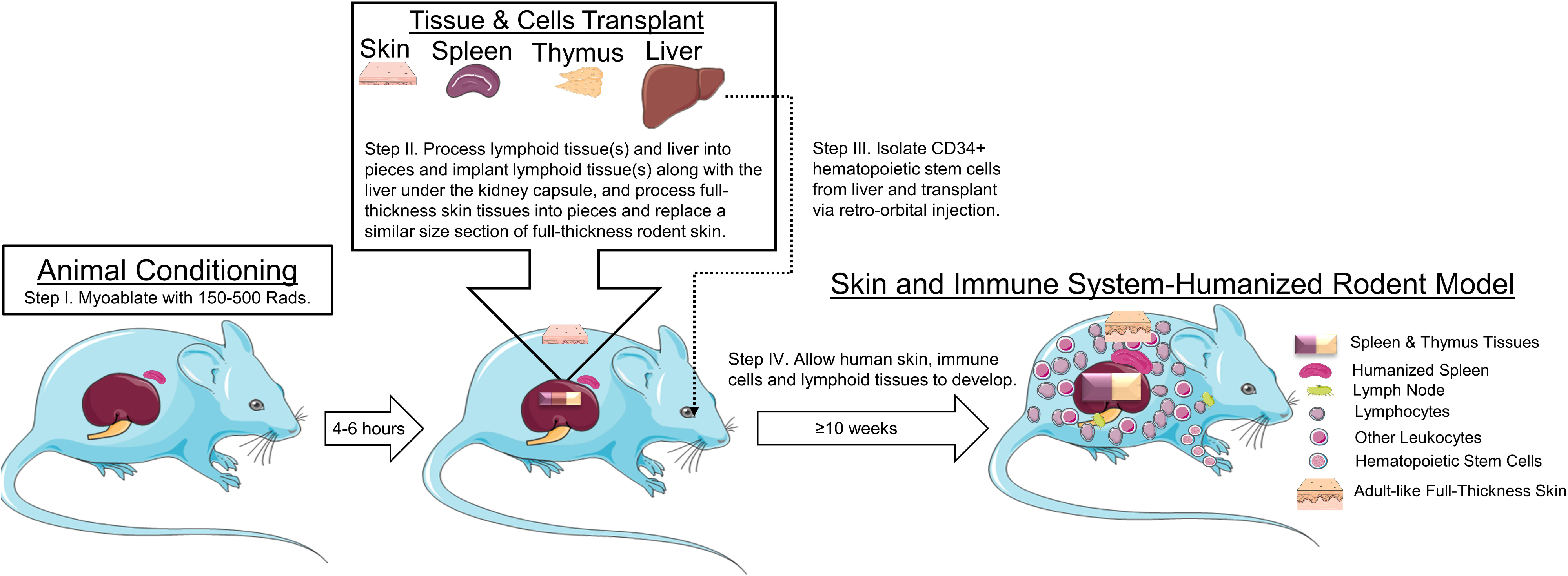
Construction of the human Skin and Immune System-humanized rodent models. (I) Severely immunodeficient rodents (mice and rats) were myoablated via gamma irradiation (150 rads-mice and 500 rads-rats) using a Cesium-137 irradiator. (II) Human fetal lymphoid tissue(s) and liver were processed into 1 mm^2^ pieces and transplanted under the renal capsule and full-thickness human fetal skin is transplanted on the panniculus carnosus of the dorsum in irradiated rodents, following the administration of antibiotics and analgesics and the induction of general anesthesia. (III) CD34^+^ hematopoietic stem cells were isolated from the fetal liver via immunomagnetic selection and transplanted via retro-orbital injection following renal capsule transplantation of the lymphoid tissues. (IV) Transplanted rodents were maintained under specific pathogen-free conditions and the human skin and lymphoid tissue(s), along with the associated immune cells were allowed to develop over a period of 10 weeks or greater, resulting in the human Skin and Immune System-humanized rodent (mouse and rat) model.

**Figure 2.**
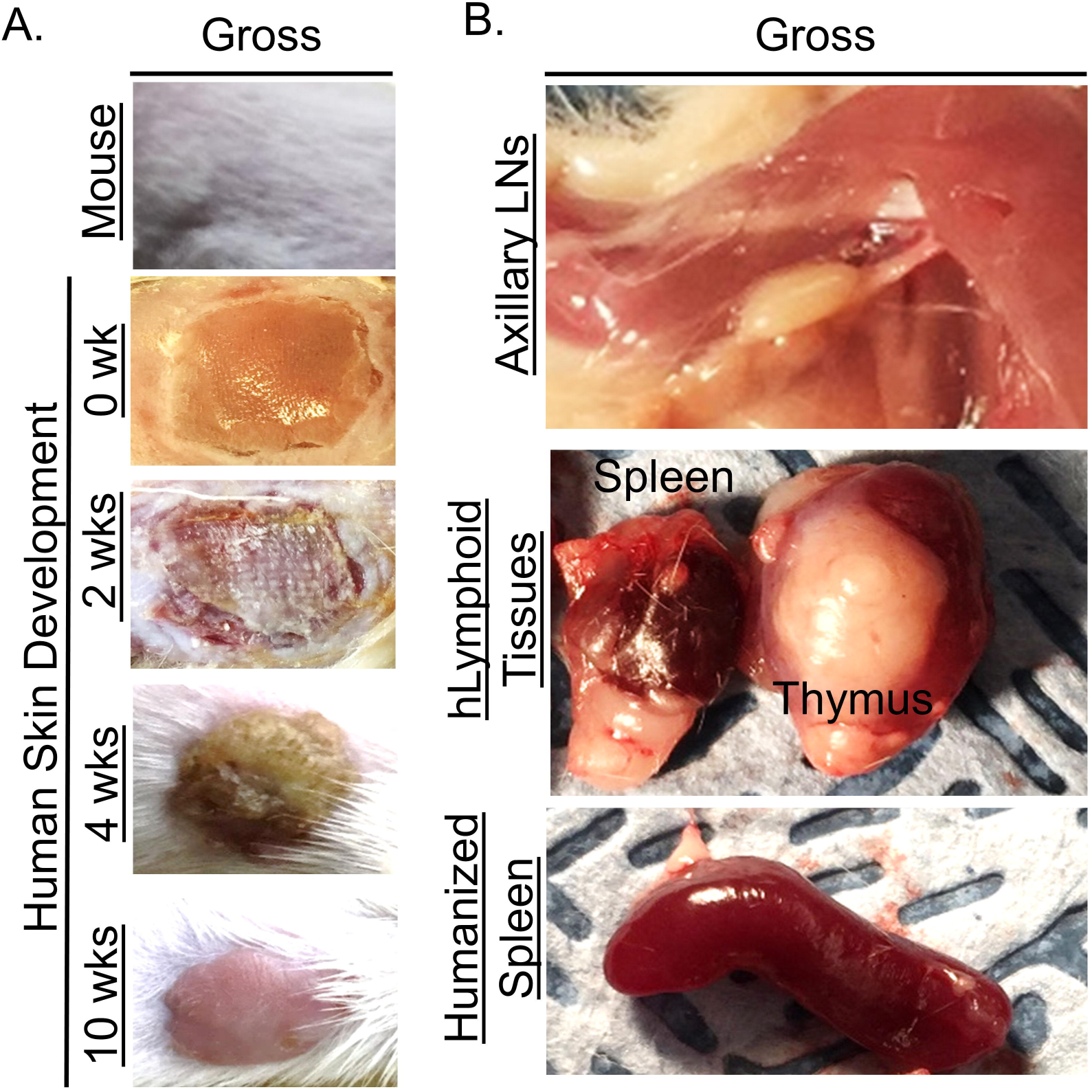
Development of human skin and lymphoid tissues in the human Skin and Immune System-humanized NSG mouse model. Transplantation of full-thickness human skin on the dorsum (A) and autologous lymphoid tissues in the kidney capsule (B) of immunodeficient mice results in robust development of full-thickness human skin and lymphoid tissues. (A) Representative gross-photos at 0 (the day of transplantation), 2, 4 and 10 weeks post-transplantation, with intact mouse skin as control. (B) Representative gross-photos of human lymphoid tissues (spleen and thymus tissues) and humanized lymphoid tissues (reconstituted-immunodeficient murine lymph node and spleen) at 10 weeks post-transplantation.

**Figure 3.**
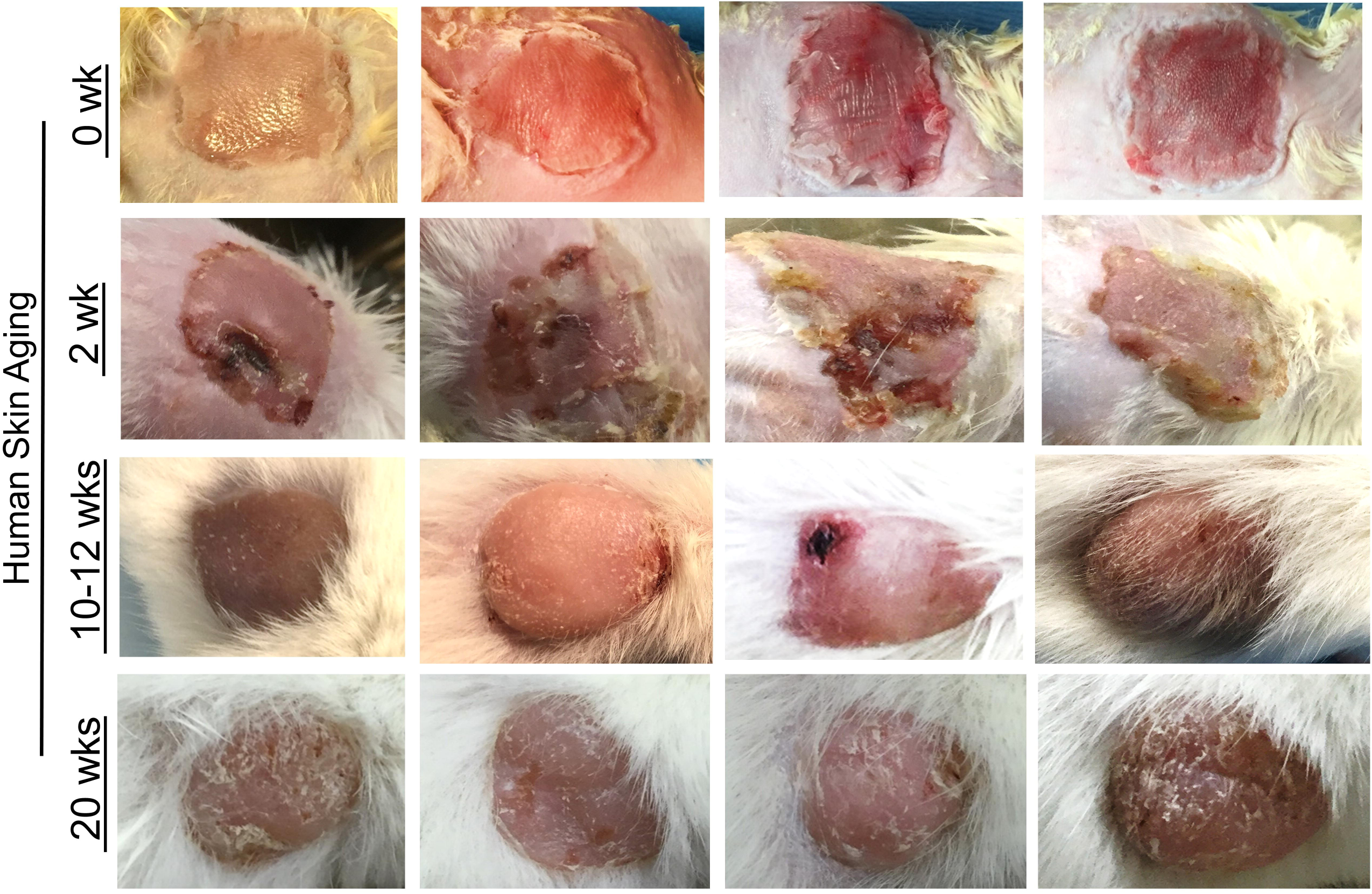
Human skin aging in the human Skin and Immune System-humanized NSG mouse model. Transplantation of full-thickness human skin on the dorsum in the hSIS-mice results in robust development of full-thickness human skin as exhibited in representative gross-photos at 0 (the day of transplantation), 2, 10-12 and 20 weeks post-transplantation; dry skin (signs of aging) and early signs of graft-versus-host disease around the human skin develops in the human skin at 20 weeks post-transplantation.

Histochemical analysis of human skin in hSIS-humanized NSG mice reveals development of the human skin graft; the microanatomy of the human skin at 10 weeks post-transplantation is comparable to adult human skin, and multiple layers of cells are present in the epidermis (**Figure 4A**). Human skin in hSIS-humanized NSG mice exhibited robust upregulation of alpha-smooth muscle actin-positive (α-SMA^+^) cells (i.e. blood vessel cells) during revascularization and wound healing (∼2 weeks post-transplantation), followed by a reduction in α-SMA^+^ cells in healed skin (∼10 weeks post-transplantation) (**Figure 4B**). The human skin exhibited robust repopulation with human keratinocytes (AE1/AE3-pan cytokeratin antibody+ cells) in epidermis and dermal fibroblasts (Anti-Fibroblasts Antibody+ cells) in the dermis (**Figure 4B**). Additionally, the human skin exhibited robust human immune cell (human CD45^+^ cells) repopulation, including Langerhans cells (CD207+ cells) (**Figure 4B**). Co-transplanted human lymphoid tissues (spleen and thymus) developed under the renal capsule (∼10 weeks post-transplantation) and exhibited microanatomy comparable to adult human lymphoid organs (**Figure 4C**) ^24^. Human thymus tissue in hSIS-humanized NSG mice exhibits robust T-cell (human CD3^+^ cells) reconstitution, with T-cell levels highest in the cortex region and relatively lower in the medulla region (**Figure 4C**) ^24,28,29^. Macrophage reconstitution (human CD68^+^ cells) in the human thymus tissue is restricted to the medulla (**Figure 4C**) ^24,29,30^. Human spleen tissue in hSIS-humanized NSG mice exhibits robust macrophage reconstitution, with macrophages predominately in the red-pulp (**Figure 4B**) ^26,27^. Additionally, the human spleen tissue in hSIS-humanized NSG mice exhibit robust T and B cell repopulation (human CD3^+^ and CD20^+^ cells), with lymphocytes predominately in the white-pulp (**Figure 4B**) ^26^. The blood plays a major role in the cutaneous immune system, as it facilitates systemic immune cell trafficking to the skin; analysis of the peripheral blood mononuclear cells (PBMCs) in hSIS-humanized NSG mice showed high (average of ∼43%) human immune cell (human CD45^+^ cells of total leukocytes) reconstitution in the peripheral blood (**Figure 5A**). Additionally, various human immune types, namely, T cells (CD4, CD8, γδ), B cells, monocytes and granulocytes were reconstituted in the peripheral blood of the hSIS-humanized NSG mouse model (**Figure 5B**).

**Figure 4.**
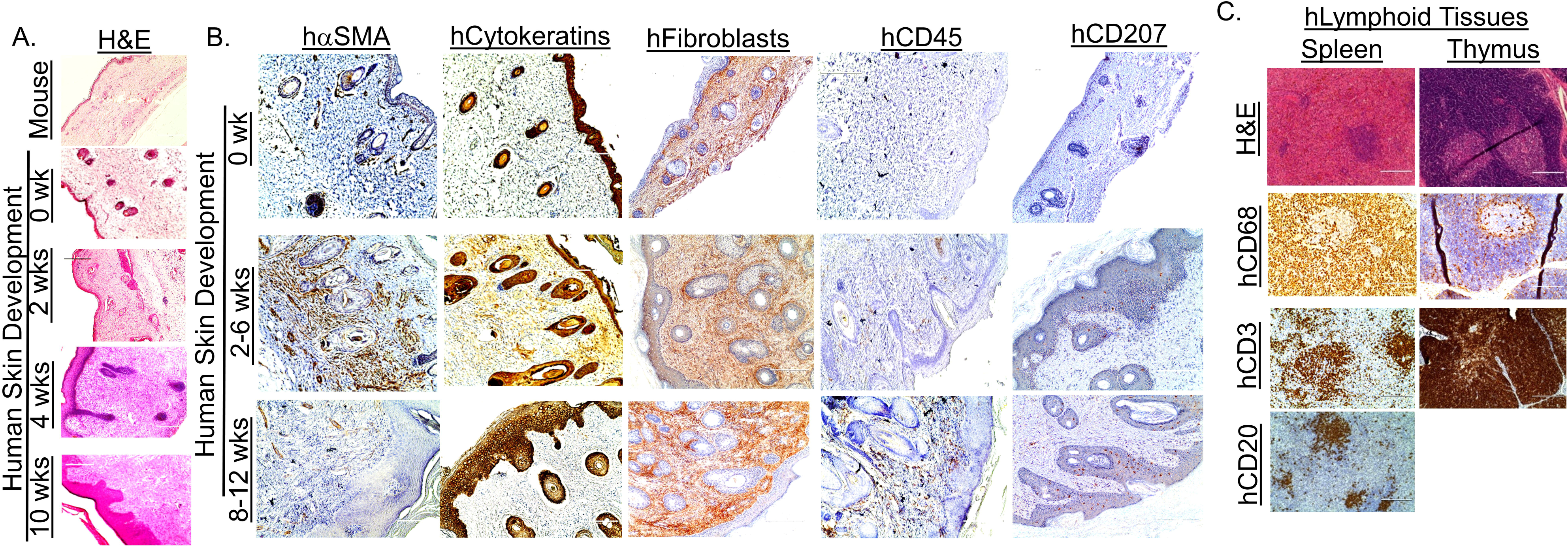
Development of human skin and immune cells in the human Skin and Immune System-humanized NSG mouse model. Representative histological (H/E) analysis of the human skin in human Skin and Immune System-humanized mice demonstrate robust development of human skin, including dermis and multicellular layer (>5 layers) epidermis and cornified envelope, which are a hallmarks of adult-human skin (A). Various human skin cells are reconstituted in the human skin, including keratinocytes (AE1/AE3+ cells, hCytokeratins+ cells), dermal fibroblast (TE7+ cells, hFibroblast+ cells), cutaneous immune cells (hCD45+ cells), and Langerhans cells (hCD207+); alpha-smooth muscle actin-expressing blood vessel cells (hα-SMA+ cells) which expands during wound healing and contrast after healing are reconstituted in the human skin. (B) Representative histological and immunohistochemical analysis of the human spleen and thymus (both under the kidney capsule) in human Skin and Immune System-humanized mice, demonstrate robust development of those tissues at 10 weeks post transplantation, with the levels of macrophages (hCD68+), T cells (hCD3+), B cells (hCD20+) comparable to human lymphoid organs. Scale bars: 200 μm.

**Figure 5.**
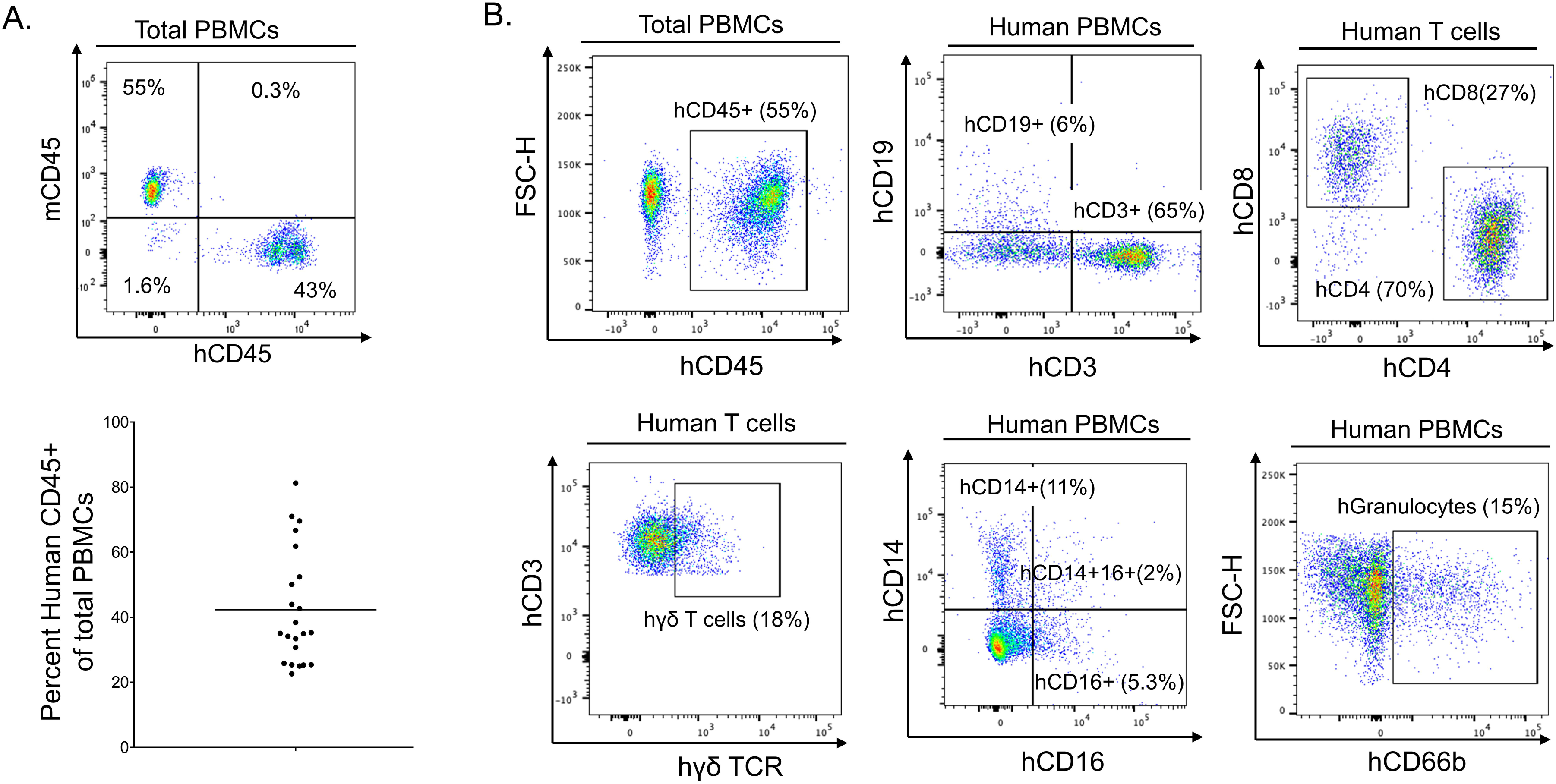
Human immune cells development in the peripheral blood in the human Skin and Immune System-humanized NSG mouse model. (A) Representative flow cytometry analysis of human immune cell (hCD45+) reconstitution in peripheral blood mononuclear cells (PBMCs) of hSIS-humanized mice at 10-12 weeks post-transplantation (A-Top Panel). Quantification of human immune cells reconstitution (n=22; 3 independent experiments) in peripheral blood mononuclear cells (PBMCs) of hSIS-humanized mice at 10 weeks post-transplantation (A-Bottom Panel). (B) A representative flow cytometry analysis of various human immune cell types (B cells-hCD19+, T cells-hCD3+, CD4+ T cells-hCD4+ T cells, CD8+ T cells-hCD8+ T cells, γδT cells-hγδTCR+ T cells, monocytes-hCD14+/hCD16+ human PBMCs, granulocytes-hCD66b+ human PBMCs) in peripheral blood mononuclear cells (PBMCs) of hSIS-humanized mice at 10-12 weeks post-transplantation.

### The hSIS-humanized SRG rat supports development of full-thickness human skin, thymus tissues, and human immune cells

Although humanized mice models have provided *in vivo* platforms for investigating the mechanisms of human diseases, the short-life span and small tissue size/volume of mice are a major limitation for pre-clinical studies. We hypothesized that a larger, severely immunodeficient rodent model with longer life span, namely rat, would support development of an *in vivo* model with longer experimental window, robust longitudinal studies (i.e. biopsy collection) and provide large tissue volume/size. We further hypothesized that co-transplantation of full-thickness fetal human skin, autologous fetal thymus and liver tissues, and fetal-liver derived hematopoietic stem cells in SRG rat would enable development of human skin, autologous thymus, and human immune cells, termed, hSIS-humanized SRG rat (**Figure 1**). We processed human thymus and liver tissues into ∼1 mm^2 pieces and isolated autologous CD34^+^ hematopoietic stem cells from the liver, and transplanted the tissues and hematopoietic stem cells into irradiated SRG rat. Full-thickness human fetal skin was processed via removal of excess fat tissues attached to the subcutaneous layer and engrafted over the rib cage where the rat skin was previously excised (**Figure 1**). Gross analysis of human skin, beginning at 3 weeks post transplantation, demonstrated robust wound healing and maturation into adult-like human skin over time (**Figure 6A**). Gross analysis of human thymus tissue development in hSIS-humanized SRG rats at 9 months post-transplantation showed robust development of the tissues under the kidney capsule (**Figure 6B**). The human skin in hSIS-humanized SRG also support the development of human skin appendages (hair) (**Supplementary Figure 2**)

**Figure 6.**
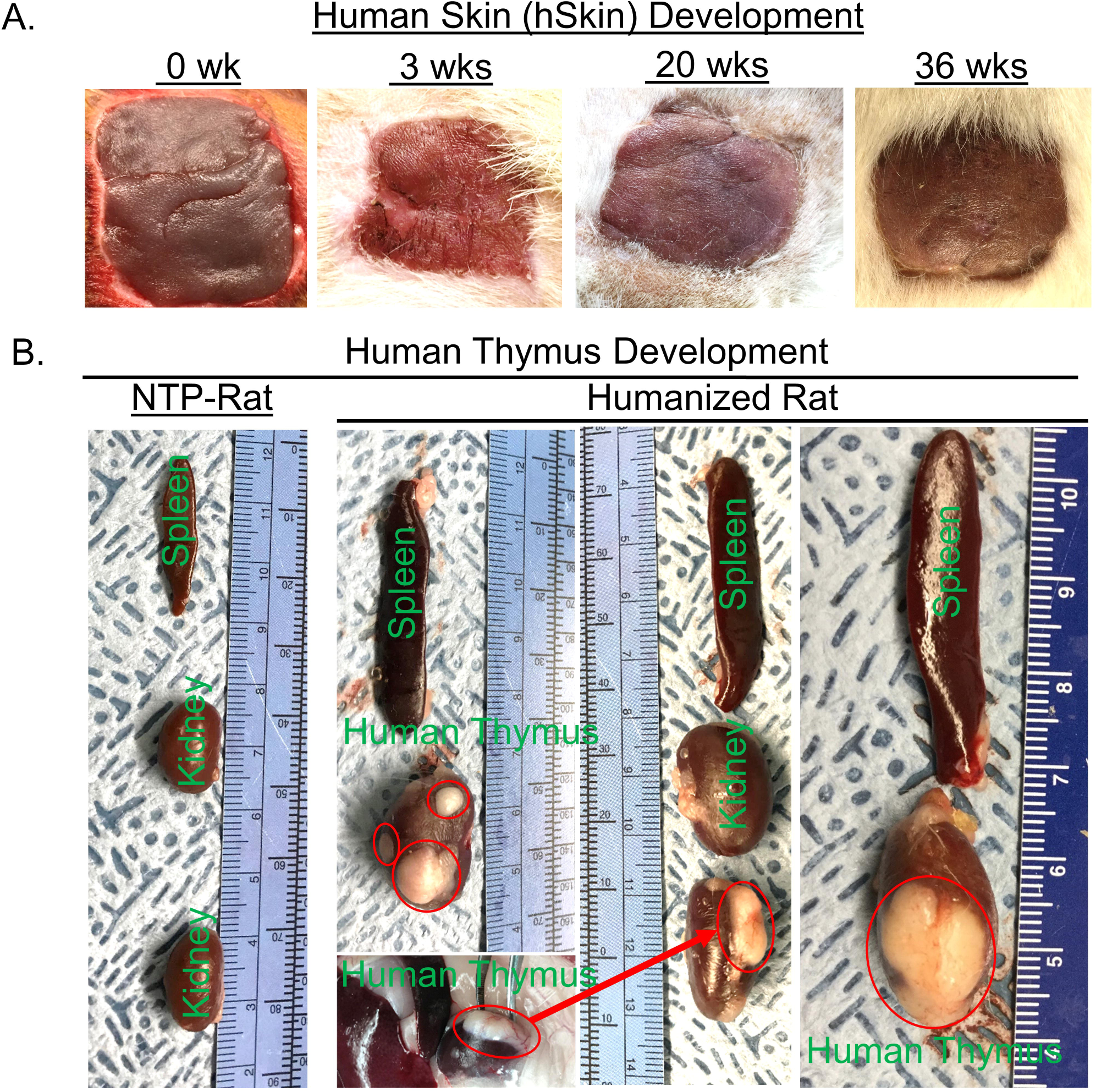
Development of human skin and lymphoid tissues in the human Skin and Immune System-humanized SRG rat model. Transplantation of full-thickness human skin on the dorsum (A) and autologous lymphoid tissue (thymus along with liver) in the kidney capsule (B) of immunodeficient rats results in robust development of full-thickness human skin and lymphoid tissues. (A) Representative gross-photos at 0 (the day of transplantation), 3, 20 and 36 weeks post-transplantation demonstrate robust human skin development. (B) Representative gross-photos of lymphoid tissues (human thymus in the kidney capsule and rat spleen) at 9 months post-transplantation demonstrates robust development of lymphoid tissues compared to non-transplanted immunodeficient rat.

Human skin in the hSIS-humanized SRG rats exhibited robust development of multi-layered human keratinocytes (AE1/AE3-pan cytokeratin antibody+ cells) in the epidermis, and dermal fibroblasts (Anti-Fibroblasts Antibody+ cells) in the dermis, both of which are comparable to adult human skin (Adult-hSkin) (**Figure 7A**). The human skin in the hSIS-humanized SRG rats exhibited robust development of human immune cells (hCD45^+^ cells), including Langerhans cells (hCD207^+^ cells), comparable to adult human skin (Adult-hSkin) (**Figure 7A**). Histochemical analysis of human thymus tissues in hSIS-humanized SRG rats at 9 months post-transplantation, exhibits a microanatomy comparable to the adult human thymus (**Figure 7B**). Human thymus tissue in the hSIS-humanized SRG rat exhibits robust human immune cell (human CD45^+^ cells) reconstitution, including T cell (human CD3^+^ cells) and macrophage (human CD68^+^ cells) reconstitution (**Figure 7B, 7C**). Additionally, human T cells in the thymus exhibit robust cytokine response to physiological stimulation using CD3/CD28 beads (**Supplementary Figure 3**). Human immune cells in the hSIS-humanized SRG rat also reconstitute the immunodeficient-rat spleen (**Figure 7D**), albeit human immune cell reconstitution of the blood was not observed (*data not shown*).

**Figure 7.**
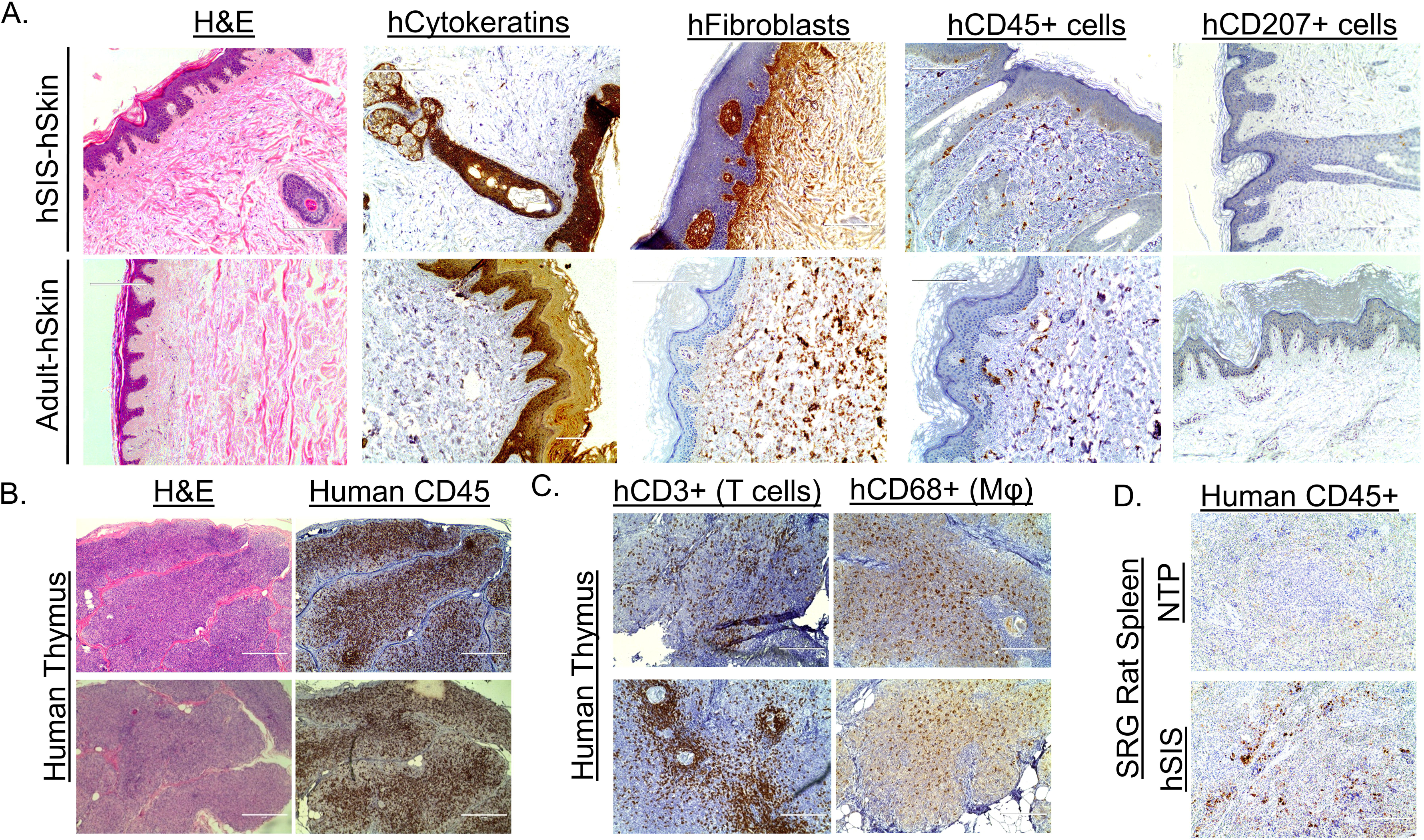
Development of human skin and immune cells in the human Skin and Immune System-humanized SRG rat model. Representative immuno-histochemical analysis of the human skin in the human Skin and Immune System-humanized rat demonstrate robust development of human skin, including dermis and multicellular layer (>5 layers) epidermis and cornified envelope, which are a hallmarks of adult-human skin (Adult-hSkin) (A). Various human skin cells are reconstituted in the human skin, including keratinocytes (AE1/AE3+ cells, hCytokeratins+ cells), dermal fibroblast (TE7+ cells, hFibroblast+ cells), cutaneous immune cells (hCD45+ cells), and Langerhans cells (hCD207+). (B) Representative histological and immuno-histochemical analysis of the human thymus (under the kidney capsule) in human Skin and Immune System-humanized rat, demonstrate robust development of human thymus tissue at 9 months post transplantation, with human immune cells (Humans CD45+), including (C) high levels of T cells (hCD3+) and macrophages (hCD68+). (D) The rat spleen in the human Skin and Immune System-humanized SRG rat model is also reconstituted with human immune cells (Humans CD45+); non-transplanted (NTP) SRG rat was used as a staining control. Scale bars: 200 μm.

### The human skin in immunodeficient rodent models supports community-associated methicillin-resistant *Staphylococcus aureus* infection and skin pathology

Community-associated methicillin-resistant *Staphylococcus aureus* (CA-MRSA) infection represents a significant public health threat ^5^; thus, understanding the host-pathogen interactions in the human skin is critical for the development of therapeutics. In order to demonstrate that human skin engrafted onto immunodeficient rodents provides a novel approach for modeling human host-skin pathogen interactions; we inoculated (intradermal inoculation) the human skin with CA-MRSA USA300. Inoculation of the human skin in immunodeficient rats with CA-MRSA results in skin lesions, comparable to lesions in CA-MRSA patients; as anticipated, no lesions was observed in non-inoculated (healthy control) human skin nor CA-MRSA-inoculated rat skin in immunodeficient rats (**Figure 8A**). CA-MRSA infection in the human skin in immunodeficient rats induced pathological changes in the epidermis and the dermis (**Figure 8B**). Concomitant CA-MRSA bacterial growth was observed in the inoculated human skin engrafted on immunodeficient rats (**Figure 8C**). Additionally, inoculation of CA-MRSA in the engrafted human skin in the NSG mouse model at 10-12 weeks post-transplantation, supports bacterial infection and skin pathology (**Supplementary Figure 4**).

**Figure 8.**
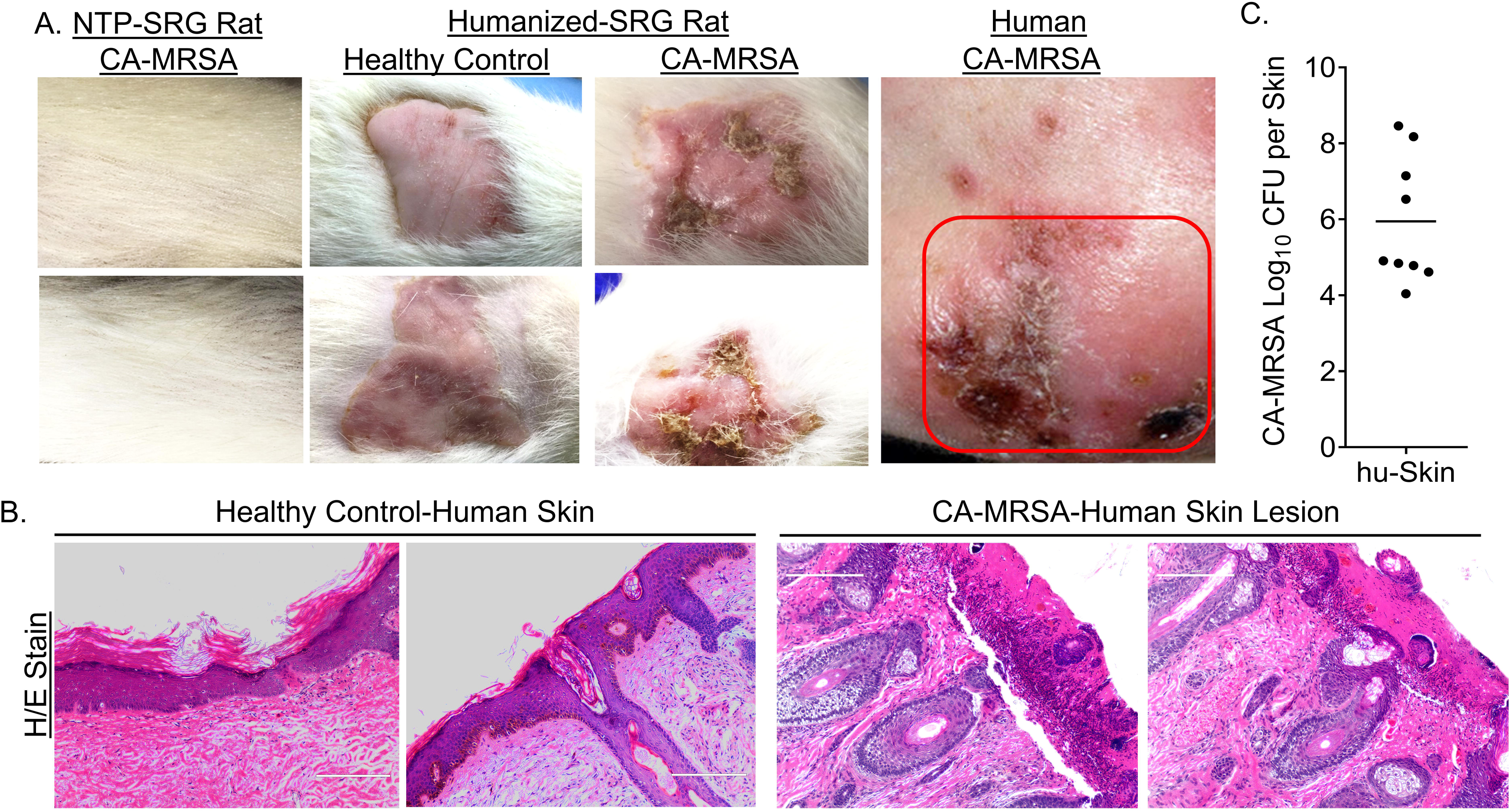
Human skin in SRG rats support CA-MRSA infection. CA-MRSA was inoculated (intradermal) into the engrafted human skin in immunodeficient rats (humanized rats) and the rat skin of non-transplanted (NTP) immunodeficient rats (NTP-Rat). (A) Gross pathology and (B) histological analysis were examined in CA-MRSA inoculated humanized rats and non-transplanted rats at 6 weeks post-inoculation, and compared to health control-human skin in humanized rats; (A) CA-MRSA infected human skin was used for comparative analysis of gross pathology in humanized rats and humans (Patient CA-MRSA skin photo credit: S. Camazine). (C) The human skin in humanized rats support high CA-MRSA bacteria load as measured at 6 weeks post-infection. Scale bars: 200 μm.

## Discussion

The human skin is the primary barrier and host defense-organ against a myriad of pathogenic microbes (i.e. CA-MRSA) and environmental insults (i.e. ultraviolet radiation) ^10^. Additionally, the skin is the primary route of infection for various emerging and re-emerging vector-borne diseases, such as tick-borne disease (i.e. Lyme disease, Rocky Mountain spotted fever, etc.) and mosquito-borne disease (i.e. Zika virus disease, Malaria, Dengue virus disease) ^7^, and also provides a robust route for vaccine and other therapeutic administration against these infectious agents ^12^. The increasing public health threat of infectious agents that target the skin (i.e. CA-MRSA) and geographic expansion of vector-borne diseases have highlighted the need for robust *in vivo* models for investigating the human skin-pathogen interactions associated with said infectious agents and evaluating therapeutics ^4-6^. Furthermore, the development of novel vaccine and therapeutic delivery platforms (i.e. skin-patch, microneedle, intradermal-vaccines) targeting the skin also illustrate the need for robust *in vivo* models with human organ systems for investigating the safety and efficacy of those vaccines and therapeutics against human infectious diseases ^12^. Rodent models are the primary platforms for investigating skin-associated infectious agents, the mechanisms of disease and host response/defense, and novel therapeutics. Although rodent models provide insights into the mechanisms of disease and host response/defense against skin-associated infectious agents, several limitations exist in rodent models, resulting in knowledge gaps. It is well established that the human skin exhibits significant structural differences compared to rodent skin ^10,25^. Specifically, human skin is composed of multiple layers and cushioned in subcutaneous fat; rodent skin, however, is composed of a single/few layers, with little to no sub-cutaneous fat ^14,25^. Differences in the types and levels of fatty acids, could play a critical role in immune signaling and regulating the skin microbiome, resulting in significant differences in anti-microbial response between human and rodent skin ^1,2^. These differences could significantly impact the translation of mechanistic findings from conventional rodent models to humans.

To address these technical gaps, larger animal models, namely pig, with skin structure and microanatomy comparable to the human skin have been employed; however, the size, cost and lack of molecular tools, limits employment of said models for mechanistic studies ^31^. An attractive alternative is transplantation of human skin and autologous immune system on immunodeficient small animal (rodent) models, namely mice and rats. Published reports demonstrated that immunodeficient mice support engraftment of human skin ^32,33^. Full-thickness human fetal skin readily engrafts onto immunodeficient mice, and develops into adult-like skin, in part due to the high regenerative capability and low MHC I and II expression, reducing the immunogenicity of the fetal skin compared to adult skin ^34,35^. Furthermore, reports demonstrate that nude rats (moderate immunodeficiency) support human skin (split-thickness skin) engraftment and development, albeit host-mediated immune rejection occurs within a few months ^36-38^. Although several rodent models demonstrate human skin engraftment and development, said platforms have not been coupled to autologous human immune cells and lymphoid tissues, such as in the hSIS-humanized SRG rodent models.

In this study, we demonstrate that co-transplanting human fetal skin along with autologous lymphoid tissues and hematopoietic stems cells in severely immunodeficient adult mice results in robust development of full-thickness, adult-like, human skin, as well as autologous human immune system cells in lymphoid tissues and the blood. In parallel, we demonstrate that co-transplanting human fetal skin along with thymus and hematopoietic stems cells in severely immunodeficient adult rat results in human skin and thymus development along with immune cells in the rat lymphoid organ (spleen), although peripheral immune reconstitution is not observed. Furthermore, major skin appendages such as hair developed on the human skin in both immunodeficient mice and rats. To establish that the human skin xenograft in immunodeficient rats is susceptible to human pathogens, we infected engrafted human skin with CA-MRSA via intradermal inoculation. We demonstrated that the human skin in immunodeficient rats supports CA-MRSA-infection and associated skin lesion, which was comparable to patients. Furthermore, the human skin in immunodeficient mice also supports CA-MRSA-infection and associated skin pathology; albeit the experimental window for infection and associated pathogenesis was restricted to less than 4 months post-transplantation, as the human skin aged rapidly (i.e. excessive dryness) and developed early sign of graft versus host disease beginning at 5 months post-transplantation. hSIS-humanized SRG rats do not exhibit rapid aging (i.e. excessive dryness) of the human skin or signs of graft versus host disease; thus providing a large experimental window (>9 months). Future efforts will improve the peripheral human immune reconstitution in immunodeficient rats.

In summary, we report the development of human Skin and Immune System-humanized (hSIS) NSG mouse and SRG rat models that incorporate human lymphoid tissues, human immune cells, and autologous human full-thickness skin. hSIS-humanized NSG mice and SRG rats provide a means of addressing several limitations in conventional rodents for modeling human skin biology/pathobiology and associated cutaneous immune responses, including microanatomy differences, susceptibility to human infectious agents and associated diseases, and cutaneous immune cell signaling. hSIS-humanized NSG mouse and SRG rat models provide robust rodent models for *in vivo* mechanistic studies of a myriad of human diseases associated with the skin, including cancer ^39,40^, viral infections ^41-50^, bacterial infections ^51,52^, etc.; thus our hSIS-humanized rodent models represents a novel *in vivo* platform for biomedical research.

## Methods

### Construction of human Skin and Immune System-humanized rodents

Adult male and female severely immunodeficient rodents (mice and rats), carry mutations in the Protein Kinase, DNA-Activated, Catalytic Subunit (PRKDC) (mice) or recombination activating gene 2 (RAG2) (rat) and interleukin-2 receptor subunit gamma (IL2RG) on the Non-obese Diabetic (NOD) strain (mice) (Jackson Laboratory, Stock No: 005557) and on the Sprague Dawley (SD) strain (rats) (Hera Biolabs) were obtained from vendor and breed in the Division of Laboratory Animal Resources (DLAR) facility at the University of Pittsburgh. Human fetal tissues were obtained from the Health Sciences Tissue Bank at the University of Pittsburgh. Human fetal tissues for constructing humanized rodents were handled and processed under biosafety level 2 conditions. Male and female rodents were myoablated via gamma radiation using cesium-137 irradiator, with mice receiving a dose of (150 rads) and rats receiving a dose of (500 rads). Myoablated male and female mice were transplanted with human fetal-thymus, liver and spleen in the kidney capsule and autologous CD34^+^ hematopoietic stem cells (via retroorbital injection of 0.2×10^6 cells) ^24^, followed by transplantation of the autologous full-thickness human fetal skin on the panniculus carnosus of the dorsum ^53-55^. Myoablated male and female rats were transplanted with human fetal-thymus and liver in the renal capsule and autologous CD34^+^ hematopoietic stem cells (via retroorbital injection of 0.5×10^6 cells), followed by transplantation of the autologous full-thickness human fetal skin (less than 4 days old) on the panniculus carnosus of the dorsum ^53-55^. In some instances rodents were only transplanted with full-thickness human fetal skin. Rodents were housed under specific-pathogen free conditions and fed irradiated chow and autoclave water.

### Immune cell reconstitution and functional assays

At indicated time points, peripheral blood was collected from animals, and leukocytes were analyzed using flow cytometry ^24^. Briefly, peripheral blood was collected from rodents and mixed with 20 mM Ethylenediaminetetraacetic acid (EDTA) at a 1:1 ratio, and single cell leukocytes were prepared via red blood cells lysis using Ammonium-Chloride-Potassium (ACK) buffer. Single-cell suspensions were stained with a LIVE/DEAD Fixable Aqua Dead Cell Stain Kit (ThermoFisher Scientific), fluorochrome-conjugated antibodies (anti-mouse CD45-BioLegend Cat. No. 103126, anti-human CD45-BioLegend Cat. No. 304014), fixed with formalin, and analyzed on a BD LSRFortessa™ cell analyzer - flow cytometer (BD Biosciences). We analyzed the data using FlowJo software (Dako). Leukocytes were selected based on forward and side scatter measurements. Single cell and live leukocytes were selected for further analysis of the percentage of human leukocytes (anti-human CD45^+^, hCD3^+^, hCD4^+^, CD8^+^, hγδTCR^+^, hCD19^+^, hCD14^+^, hCD16^+^, hCD66b^+^) and mouse leukocytes (anti-mouse CD45^+^). The analysis of the various human immune cell populations and subsets were gated on human leukocytes. Human T cells were also isolated from the thymus tissue in humanized SRG rats via immunomagnetic selection using anti-human CD3 antibody (EasySep™ Human CD3 Positive Selection, Catalog # 17951, Stemcell Technologies) and treated without (vehicle) or with Gibco™ Dynabeads™ Human T-Activator CD3/CD28 (Cat. No. 111.61D, ThermoFisher Scientific) in the presence of recombinant IL2 and BD GolgiPlug (BD Biosciences) for 12 hours. Human cytokines expression (hTNFα and hIFNγ) in human T cells were analyzed using BD LSRFortessa™ cell analyzer - flow cytometer (BD Biosciences) and the data were analyzed using FlowJo software (Dako).

### Gross/In situ immune cell analysis

Gross analysis of tissues was performed using a camera (8 mega pixel), with animals either euthanized or anesthetized prior to photographing. Indicated tissue samples from humanized rodents or humans (adult human skin, BioChain, catalog number: T2234218), were fixed with formalin, and subsequently embedded in paraffin. Paraffin embedded fixed sections were stained via hematoxylin and eosin or with indicated human antibodies ^24^ (anti-human CD45-Biocare Medical Cat. No. CME PM016AA; anti-human CD3-Biocare Medical Cat. No. CME 324 A, B, C; anti-human CD68-Biocare Medical catalog number CM 033 A, B, C; anti-human CD20-Biocare Medical catalog number ACR 3004 A, B; anti-human alpha smooth muscle actin; anti-pan cytokeratin, Clone AE1/AE3, Biocare Medical catalog number SKU: 011; anti-human fibroblast, Clone TE7, Millipore Sigma catalog number CBL271; anti-human CD207, Dendritics catalog number: DDX0362). The immunoreactivity of the antibodies were determined via incubation with DAB substrate (MACH 2 Detection Kit, Biocare Medical) and counterstaining with hematoxylin.

### CA-MRSA infection in the human skin in immunodeficient rodents

The human skin on immunodeficient rodents were inoculated with CA-MRSA USA300 ^56^ via intradermal injection with 1×10^8 for rats and 1×10^6 for mice; non-transplanted immunodeficient rats and mice were inoculated with the same dose for controls. Portions of equal weight of human or rodent skin was excised and bacterial load based on the number of colony forming units (CFU); CA-MRSA bacteria strain was confirmed via polymerase chain reaction (PCR).

### Study approval

De-identified human fetal tissues at the gestational age of 18 to 20 weeks were obtained from medically or elective indicated termination of pregnancy through Magee-Women’s Hospital of the University of Pittsburgh Medical Center (UPMC), with the University of Pittsburgh, Health Sciences Tissue Bank. Written informed consent of the maternal donors were obtained in all cases, under a protocol reviewed and approved by the Institutional Review Board (IRB) of the University of Pittsburgh; approved guidelines and federal/state regulations were adhered to for all procedures. The use of de-identified human fetal tissues to construct humanized rodents was reviewed and approved by the University of Pittsburgh’s IRB Office. The use of de-identified human fetal tissues did not constitute human subjects research as defined under federal regulations [45 CFR 46.102(d or f) and 21 CFR 56.102(c), (e), and (l)]. The use of human fetal liver-derived hematopoietic stem cells was reviewed and approved by the Human Stem Cell Research Oversight (hSCRO) at the University of Pittsburgh. The use of a biological agent (CA-MRSA), and recombinant DNA and transgenic animals was reviewed and approved by the Institutional Biosafety Committee (IBC) at the University of Pittsburgh. All animal studies/experimental protocols were reviewed and approved by the Institutional Animal Care and Use Committee at the University of Pittsburgh and were conducted following approved guidelines, which adheres to the NIH guidelines for housing and care of laboratory animals.

## Acknowledgments

We used the UPMC-Hillman Cancer Center and Tissue and Research Pathology within the University of Pittsburgh’s Biospecimen Core, which is supported in part by the NIH award P30CA047904. Berthony Deslouches, Department of Microbiology and Molecular Genetics; University of Pittsburgh provided insightful advice on developing this project. This work was supported by the National Institutes of Health (NIH)-National Institute of Allergy and Infectious Diseases (NIAID) (R21AI135412) and National Institutes of Health (NIH)-Fogarty International Center (FIC) (D43TW010039). Diagrams and experimental schemes were made via modification of images from Creative Commons.

## Authors contributions

MB, LT, TY and AR conceived and designed experiments in the study. YA, CB, SH, AD, LT, RS, SK, SB, IC and MB performed experiments. MB, AR, LT, YA and CB analyzed and interpreted the data. MB, YA, CB and SH prepared the manuscript.

## Competing interests

Yash Agarwal, Cole Beatty, Sara Ho, Lance Thurlow, Antu Das, Samantha Kelly, Rajeev Salunke, Isabella Castronova, Shivkumar Biradar, Anthony Richardson, and Moses Bility have declared that no conflict of interest exists. Tseten Yeshi has a financial conflict of interest, as he works for HaraBiolabs, which provides the Sprague-Dawley-Rag2^tm2hera^ Il2rg^tm1hera^ (SRG) rat commercially.

