## Supplementary Figures for "Development of humanized mouse and rat models with full-thickness human skin and autologous immune cells"

**Supplementary Figures and Figure Legends**

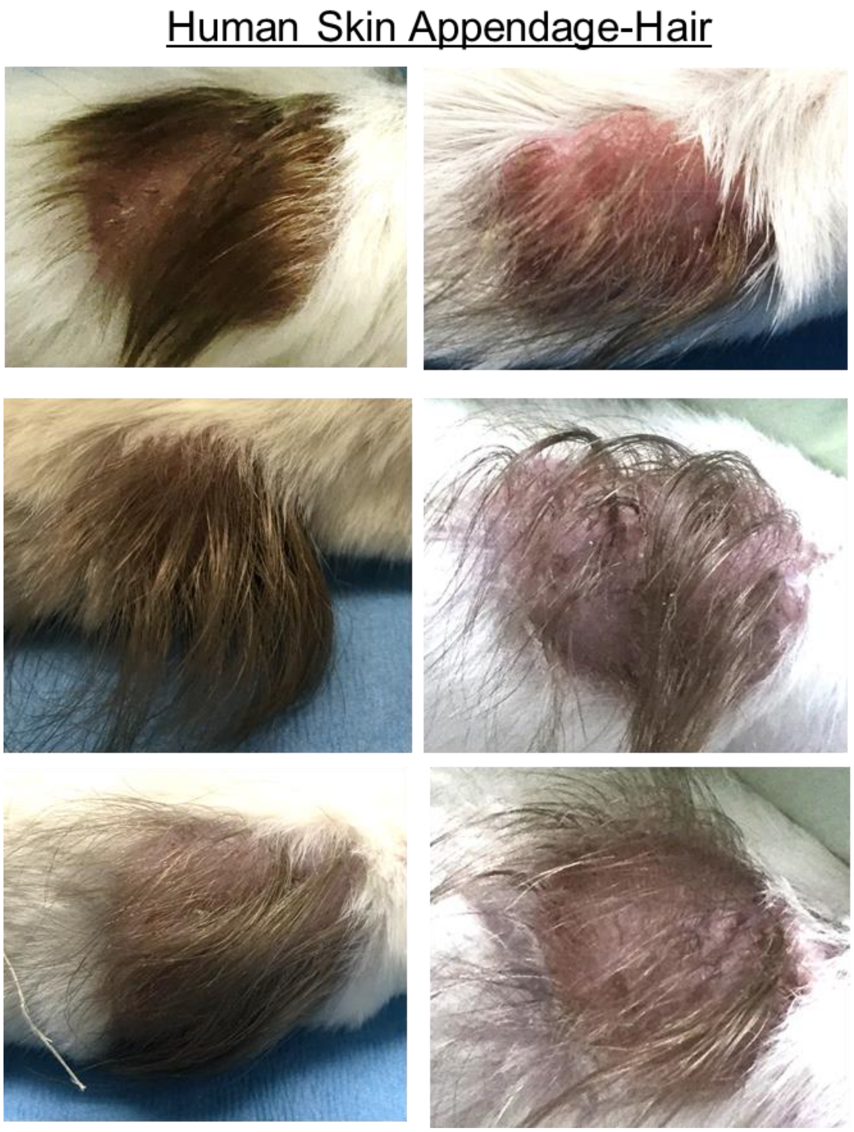


**Supplementary Figure 1. Development of human skin appendage-hair in the human Skin and Immune System-humanized NSG mouse model.** Transplantation of full-thickness human skin from regions with significant hair follicles in hSIS-mice results in robust development human hair as exhibited in representative gross-photos at 12 weeks post-transplantation.

### Human Skin Appendage-Hair

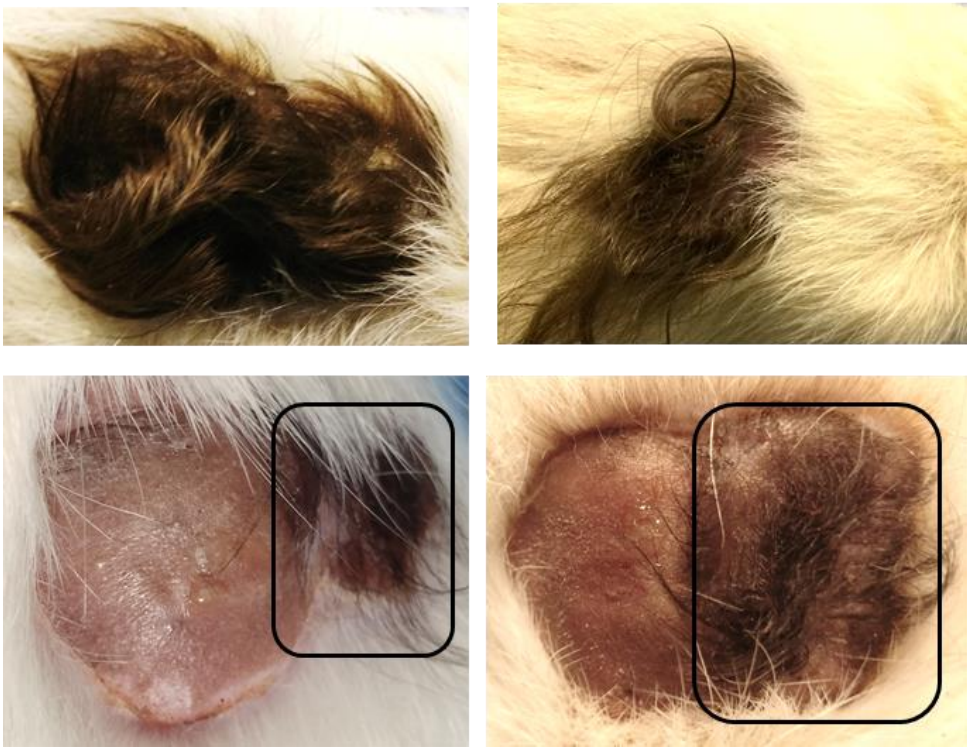

**Supplementary Figure 2. Human hair development in engrafted human skin in SRG rats.** Transplantation of full-thickness human skin from regions with significant hair follicles in hSIS-rats results in robust development human hair as exhibited in representative gross-photos at 6 months post-transplantation. In the bottom panel, skin from regions with (Right-Black box) or without (Left) significant hair follicles from the same donor were co-transplanted (adjacent to each other) to demonstrate the restriction of human hair development to regions with preexisting-hair follicles.

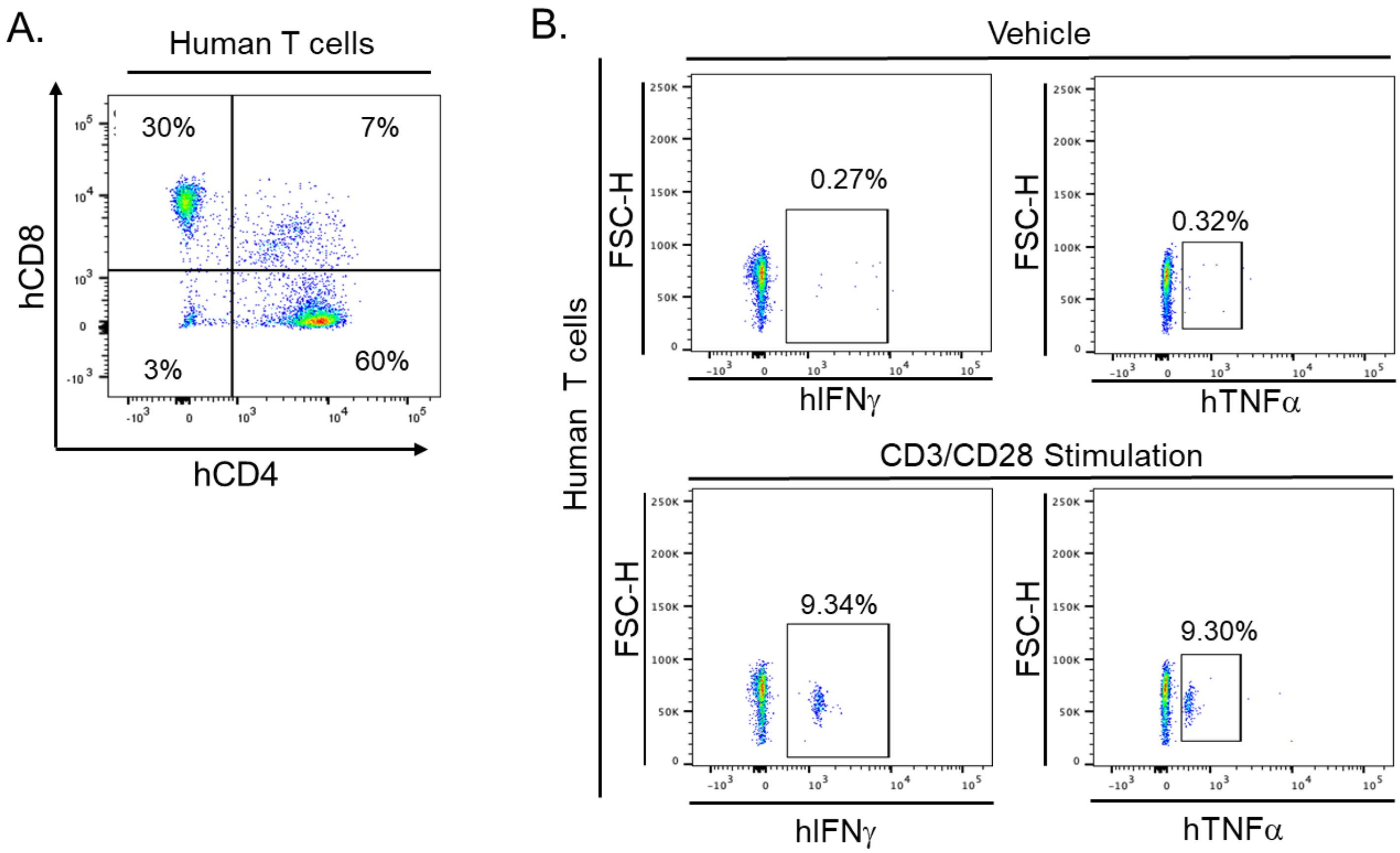


**Supplementary Figure 3. Human Skin and Immune System (hSIS)-humanized SRG rat model supports the development of functional human T cells.** (A) Representative flow cytometry analysis of human T cells (hCD3+ cells) from the human thymus tissue of hSIS-humanized SRG rat at 36 weeks post-transplantation. (B). Flow cytometry analysis of cytokine response in human T cells from human thymus tissue following stimulation without (Vehicle) or with CD3/CD28 beads.

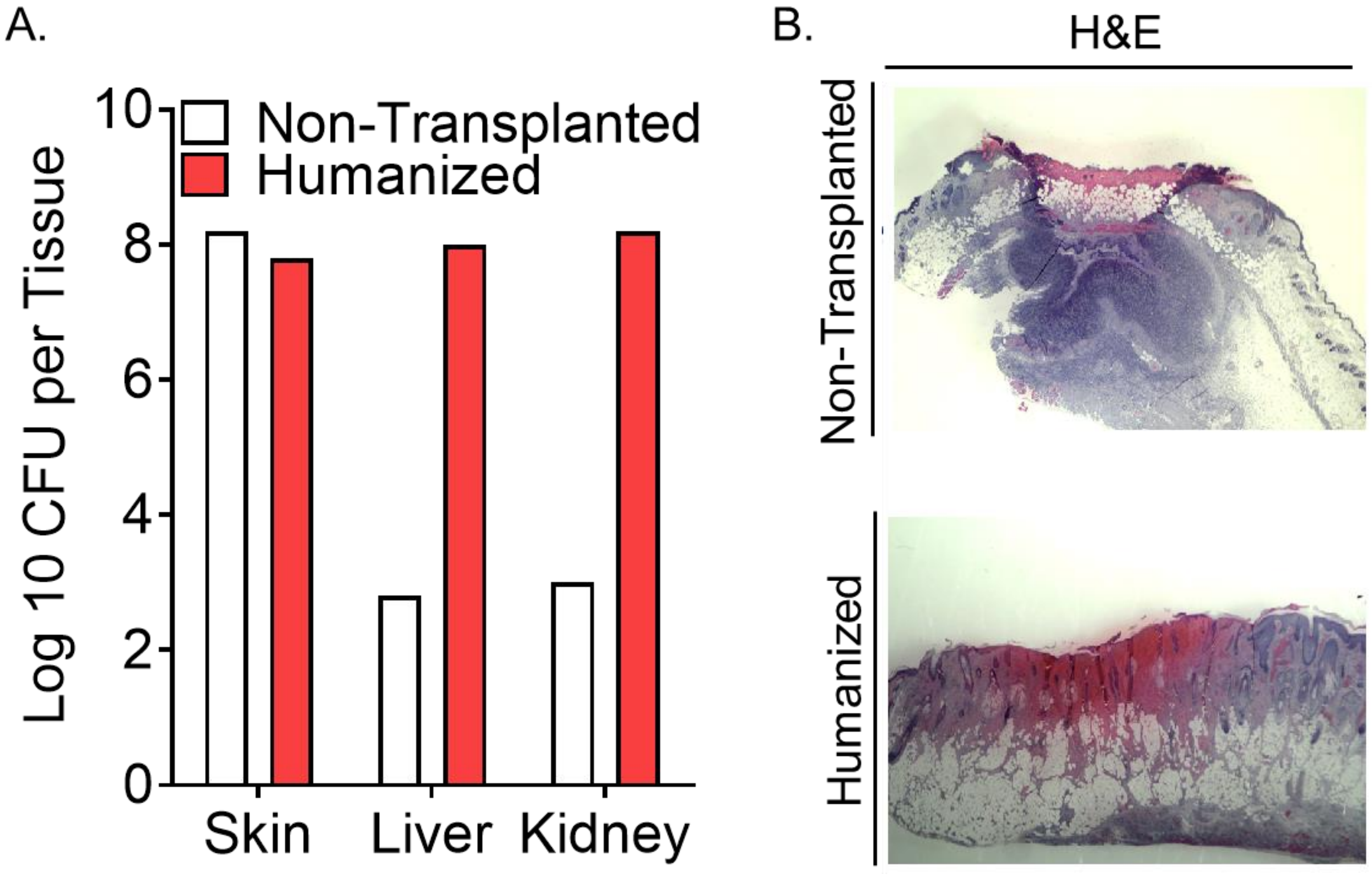


**Supplementary Figure 4. Human skin xenograft in NSG mouse model support CA-MRSA infection**. CA-MRSA was inoculated (intradermal) into the engrafted human skin in hSIS-humanized NSG mouse (Humanized) and the mouse skin of non-transplanted NSG mouse (Non-transplanted). (A) Bacterial load in the inoculated skin tissues and other tissues were examined at 3 days post-inoculation. (B) Skin pathology was examined in the inoculated skin tissues via histological (Haemotoxylin and Eosin-H&E) analysis at 3 days post-inoculation.
